# Quantum-Classical Reservoir Computing to Predict Influenza A/H3N2 Antigenic Distance

**DOI:** 10.64898/2026.07.21.739885

**Authors:** Mehdi Khalaj, Lingling Jin, Steven Rayan

## Abstract

Accurate prediction of antigenic distance between influenza A/H3N2 strains is essential for timely vaccine strain selection, yet traditional hemagglutination inhibition (HI) assays are labour-intensive and limited in throughput. We present FluQRC, a hybrid Quantum-Classical Reservoir Computing framework for sequence-based antigenic distance prediction. FluQRC integrates three novel components: (1) a differentiable gated property ranking network for data-driven property selection, (2) a dimensionality reduction network that compresses the feature representation into a form suitable for quantum processing, and (3) a hybrid quantum-classical reservoir computing architecture for antigenic distance prediction. Experiments on two datasets covering 1963–2002 (271 strains, 73,441 pairs) and 2003–2025 (888 strains, 788,544 pairs) show that FluQRC outperforms four established baselines across all three evaluation metrics (MAE, RMSE, *R*^2^). On the larger and more challenging 2003–2025 dataset, FluQRC achieves MAE = 0.369, RMSE = 0.635, and *R*^2^ = 0.900, corresponding to a 20.3% reduction in MAE and a 14.3% reduction in RMSE relative to the strongest baseline, while raising *R*^2^ from 0.862 to 0.900. These results demonstrate the scalability and effectiveness of FluQRC for large-scale antigenic distance prediction.

## I. Introduction

Influenza A virus is an enveloped virus with a negative-sense single-stranded RNA genome. Among the various influenza A subtypes infecting humans, H1N1 and H3N2 are the dominant strains responsible for seasonal outbreaks, which remains a major public health concern. Global estimates suggest that influenza viruses infect nearly one billion individuals every year, with approximately 3–5 million cases progressing to severe illness and hundreds of thousands of deaths occurring annually [1–4]. Because of this substantial disease burden, effective surveillance and vaccine design strategies are essential for controlling the spread of influenza viruses.

Vaccination is widely recognized as the most effective intervention for reducing influenza-related morbidity and mortality [5]. The hemagglutinin (HA) protein plays a central role in viral infection and immune recognition. It mediates the attachment of viral particles to host cells by binding to sialic acid receptors and facilitates membrane fusion during viral entry [6–8]. Because of its essential biological function and strong immunogenicity, HA is the primary target of neutralizing antibodies and therefore serves as the main antigenic component in influenza vaccines [9]. However, influenza viruses evolve rapidly. Over time, the mutations gradually alter the antigenic characteristics of the virus in a process known as antigenic drift [10, 11]. This evolutionary mechanism allows influenza viruses to evade previously acquired immunity and continue circulating in human populations. It is particularly evident in the A/H3N2 subtype, which evolves more rapidly than other seasonal influenza viruses and frequently undergoes transitions between antigenic clusters [12]. Historical analyses have shown that H3N2 viruses evolved through multiple antigenic clusters over time. Because of this continuous antigenic evolution, influenza vaccines must be periodically updated to match circulating viral strains.

Researchers currently rely heavily on the hemagglutination inhibition (HI) assay to measure antigenic similarity between viral strains and has become the standard laboratory technique for antigenic characterization and estimation of antigenic distance between circulating influenza viruses and reference vaccine strains [13]. Despite its reliability, the HI assay involves extensive laboratory procedures and provides only moderate throughput, which limits its applicability for largescale or rapid screening of emerging viral variants [14]. With the rapid progress of sequencing technologies and the growing availability of viral genomic sequences and serological datasets, researchers have increasingly explored computational approaches to infer viral antigenicity [15]. Such methods enable rapid, scalable analysis of viral evolution and serve as valuable complements to traditional serological assays. Predicting antigenic distance from sequence data is a core task in antigenicity prediction, as it directly determines whether a newly emerged strain is sufficiently different from the current vaccine strain to be classified as an antigenic variant requiring a vaccine update.

To estimate antigenic differences, several studies formulated the problem as a regression task in which antigenic distance is predicted from sequence data. Earlier investigations had shown that antigenic changes are typically driven by mutations at a relatively small number of critical residues [16]. Building on this observation, regression-based models such as support vector regression and joint random forest regression were proposed to establish quantitative relationships between HA sequence variation and antigenic distances [17, 18]. With the rapid development of machine learning and deep learning, more sophisticated predictive models have been introduced. Early efforts applied nonlinear regression [19] and convolutional neural networks [20] to estimate antigenic distances directly from HA sequences. Subsequent work refined this paradigm with contrastive learning to capture subtle sequence variations [21], BiLSTM-based language-model embeddings combined with ridge regression to reconstruct antigenic maps [22], and graph neural networks coupled with meta-learning to transfer knowledge across influenza subtypes under limited labeled data [23]. Despite their strong performance, most sequence-based models overlook the physicochemical properties of amino acids, even though factors such as antigenic regions and HA glycosylation are known to drive antigenicity. To address this, Liao et al. [24] grouped amino acids by polarity, charge, hydrophobicity, and volume, and modelled antigenic relationships through regression on HA mutations. More recent frameworks integrate multiple biological descriptors: MFPAD [25], for instance, combines substitution counts, glycosylation site differences, and physicochemical property differences within an XGBoost regressor to predict antigenic distances.

Although these approaches improve predictive performance, many depend heavily on manually designed features. Constructing such features often requires extensive biological knowledge and may limit the scalability and generalization of predictive models. To overcome this limitation, some studies have attempted to automate feature extraction using large physicochemical property databases. Cui et al. [26] identified 18 antigenically important residues in the HA1 region of A/H3N2. Mutations at these sites were encoded as physicochemical differences derived from clustered AAIndex1 properties and fed into a multiple linear regression model. Similarly, Yao et al. [18] proposed a joint random forest regression method that incorporated substitution matrices derived from physicochemical properties stored in the AAIndex2 database. Geng et al. [27] proposed FluAttn, an attention-based framework for predicting A/H3N2 antigenicity. It mines physicochemical descriptors and substitution matrices from the AAIndex database (https://www.genome.jp/aaindex/) using an attention mechanism to rank feature importance, then trains a multilayer perceptron to estimate antigenic distances between strains.

Although many classical machine learning and deep learning models have been proposed for this task, these approaches often depend on manually designed features or struggle to capture the complex, nonlinear relationships between amino acid substitutions and the resulting changes in antigenic distance. A key underlying challenge is that the mapping from sequence mutations to antigenic change is high-dimensional and governed by subtle interactions among multiple amino acid properties, which are difficult for classical models to represent efficiently. Recent advances in quantum computing offer a promising approach for addressing this challenge. Quantum systems can naturally represent high-dimensional, nonlinear mappings through quantum superposition and entanglement. These properties are inherently suited to modeling the complex feature interactions that classical approaches struggle to capture. In particular, Quantum Reservoir Computing (QRC) exploits the rich dynamics of quantum systems to project input data into an exponentially large Hilbert space, enabling powerful nonlinear representations without the need for explicit optimization of the reservoir parameters. This study investigates the application of a Hybrid Quantum-Classical Reservoir Computing technique to predict antigenic distance between influenza A/H3N2 strains, aiming to improve computational antigenicity prediction.

The main contributions of this work include: (1) a Differentiable Gated Property Ranking Network for data-driven property selection, (2) a Dimensionality Reduction network that compresses the feature representation into a form suitable for quantum processing, and (3) a hybrid Quantum-Classical Reservoir Computing architecture for antigenic distance regression.

## II. Preliminaries

This section briefly introduces the two computational paradigms that underpin our framework: classical Reservoir Computing and its quantum extension, Quantum Reservoir Computing. A full mathematical treatment of both paradigms is provided in Appendix A, B.

### A. Reservoir Computing

Reservoir Computing (RC) [28] is a supervised learning framework in which an input signal is projected into a high-dimensional dynamical space through a fixed, randomly initialised nonlinear system called the *reservoir* (Fig. 1). The reservoir produces rich internal representations of the input, from which a simple linear readout layer learns the desired mapping. Crucially, only the readout weights are trained, while the reservoir parameters remain frozen, which avoids costly gradient-based optimisation of the recurrent dynamics and yields a closed-form, ridge-regression solution for the readout. The most common digital implementation is the Echo State Network (ESN), in which the reservoir is realised as a sparsely connected recurrent neural network satisfying the Echo State Property, ensuring fading memory of past inputs.

**Fig. 1.**
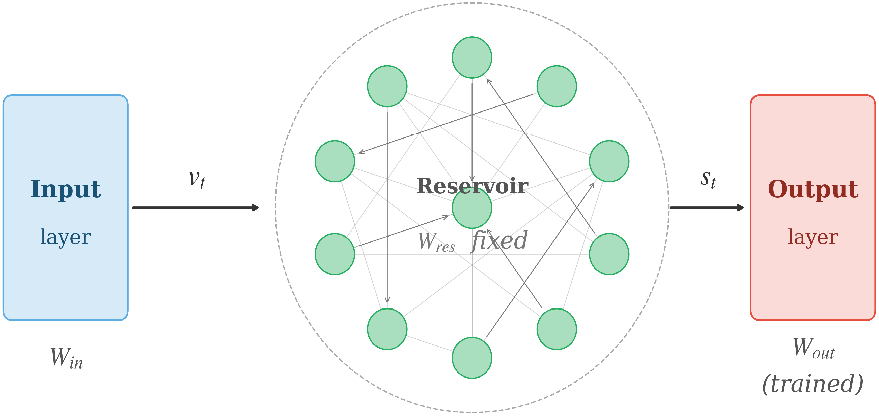
RC framework with ring-topology reservoir: sparsely connected reservoir nodes maintain fixed random weights *W*_res_, while only the readout weights *W*_out_ are trained.

### B. Quantum Reservoir Computing

Quantum Reservoir Computing (QRC) [29, 30] replaces the classical reservoir with a quantum system, exploiting properties such as superposition, entanglement, and the exponentially large Hilbert space to obtain a far more expressive feature mapping than is feasible classically. A QRC pipeline consists of four stages: (i) encoding classical inputs into quantum states, (ii) evolving these states under a fixed unitary that defines the reservoir dynamics, (iii) measuring expectation values of selected observables to extract classical features, and (iv) feeding these features into a simple classical readout for training and prediction. Because only the readout is trained, QRC inherits the efficiency of classical RC while sidestepping the barren-plateau issue that hampers gradient-based variational quantum algorithms [31].

Although QRC was initially developed and widely applied for time-series analysis, its applicability is not limited to temporal data. Recent studies have shown that QRC can be effectively extended to a broader range of problems, including applications in quantum chemistry simulations and hybrid quantum–classical neural network design [32, 33]. These developments highlight that QRC serves as a versatile computational paradigm capable of addressing diverse machine learning and scientific computing tasks beyond its original scope. Since antigenic distance prediction is not a time-series problem, we do not rely on the temporal memory of the reservoir; instead, we exploit its nonlinear, high-dimensional feature mapping to encode pairwise amino acid descriptors into representations suitable for regressing antigenic distances between influenza A/H3N2 strains.

## III. Methodology

This section presents the FluQRC framework in detail, which includes the data collection process, selecting and weighting antigenicity-relevant amino acid properties, the construction of input feature vectors from the selected properties and sequence pairs, and compressing these features for quantum processing. Finally, the hybrid quantum–classical reservoir computing architecture that predicts antigenic distances from the resulting representations is described.

### A. Data Collection

The dataset in this study consists of three main components:

1. HA sequences of influenza A/H3N2 viruses collected from the Global Initiative on Sharing All Influenza Data (GISAID)^1^ and the NCBI Influenza Virus Resource^2^. Because the HA1 subunit contains major antigenic sites, the analysis focused on the HA1 region, comprising approximately 330 amino acid positions per strain after preprocessing, 2) amino acid descriptor including **scalar physicochemical properties** and **amino acid substitution matrices** from the AAindex database, as well as Kidera factors, Z-scale descriptors, VHSE descriptors, Atchley factors, and ProtParam-derived physicochemical properties, along with substitution matrices from the PAM and BLOSUM families, and 3) experimentally validated serological measurements from the seminal work of Smith et al. [16], containing 4,228 HI titers collected between 1968 and 2002, and from annual and interim reports of the Worldwide Influenza Centre (WIC), encompassing 15,077 HI measurements spanning the period from 2003 to 2025.

Pairwise antigenic distances were constructed from the cleaned HI titer matrix using antigenic cartography. Specifically, the strain-by-strain HI matrix was embedded into a twodimensional antigenic map via the multidimensional scaling implementation in Racmacs [34], which positions each strain such that map distances best preserve the antigenic relationships implied by the HI titers. The pairwise antigenic distance was then computed as the Euclidean distance between the optimized coordinates of each strain pair, and these distances served as the ground-truth labels for training and evaluating the prediction model.

### B. Property Selection via Differentiable Gated Ranking

The goal of this section is to identify, from a large pool of heterogeneous amino acid descriptors, a compact and informative subset whose weighted combination best supports antigenic distance prediction. We proceed in six steps:

#### 1) Amino Acid Property Descriptor

To represent the biochemical differences between two viral sequences as structured numerical features, each amino acid property is encoded as a site-level descriptor of a sequence pair. Let *N* denote the total number of viral sequences, each aligned to *L* non-conserved positions. Multiple sequence alignment was performed using MUSCLE [35]. As a quality-control step, HA1 sequences containing one or more gap characters (a total of 3 sequences) were discarded. This ensured that all retained sequences were gap-free and directly comparable across aligned positions during pairwise feature extraction.

The HA1 sequence at the non-conserved positions for a virus strain *v*_*i*_ is denoted by 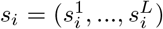, where L is the sequence length and each residue 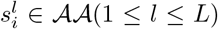. The standard amino acid alphabet is defined as:

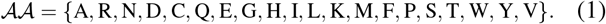

We follow the procedure described in [27] to calculate the site-level descriptor at position *l*. The collection of amino acid properties used in this work is denoted *P* . Each property *p* ∈ *P* belongs to one of two categories, with no overlapping between them:

- **Scalar physicochemical index**. A mapping *ϕ*_*p*_ : *AA* → ℝ that assigns a single real value to each amino acid. For a sequence pair (*s*_*i*_, *s*_*j*_), the site-level descriptor at position *l* is the absolute difference of property values:

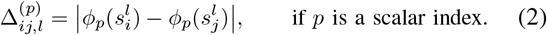
- **Substitution matrix**. A mapping *M*_*p*_ : *AA* × *AA* → ℝ that assigns a score to each ordered pair of amino acids. For a sequence pair (*s*_*i*_, *s*_*j*_), the site-level descriptor at position *l* is the substitution score directly:

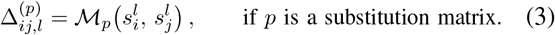

Collecting the site-level descriptors across all *L* positions yields a property-specific descriptor vector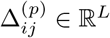 . Stacking descriptors for all *P* = |*P*| properties column-wise yields the descriptor matrix for the sequence pair (*s*_*i*_, *s*_*j*_):

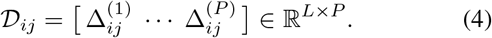

This matrix provides a unified *L* × *P* representation of pairwise amino acid variation across all properties and positions.

#### 2) Property Pre-processing

Before descriptor construction, two pre-processing steps are applied to ensure scale uniformity across heterogeneous descriptor sources before learning. For scalar indices, we removed redundant properties using a Pearson correlation filter, retaining only one property from any pair with an absolute correlation above *ρ*_max_ = 0.6. The remaining scalar properties are then standardized to zero mean and unit variance across the twenty standard amino acids. For substitution matrices, asymmetric ones were excluded to ensure consistency in pairwise amino acid comparisons. The entries of all selected features were subsequently range-normalized to the interval [0, 1].

#### 3) Differentiable Gated Property Ranking Network

The central goal of property selection is to identify, from the full set *P* of descriptors, a compact subset *P*^*′*^ of size *K* that is most informative for predicting antigenic distance. To this end, we propose the *Differentiable Property Ranking Network* (DPRN), in which property importance is encoded as a set of learnable gates 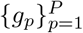 , with each *g*_*p*_ ∈ (0, 1) obtained by passing a learnable logit through a temperature-controlled sigmoid (Fig. 2). This design follows the principle of differentiable feature selection via continuous relaxation of discrete gates [36]. Crucially, each gate *g*_*p*_ is independent — unlike softmaxm normalisation, which forces competition among properties — allowing multiple properties to simultaneously receive high importance scores. The full formulations of DPRN are provided in Appendix C.

**Fig. 2.**
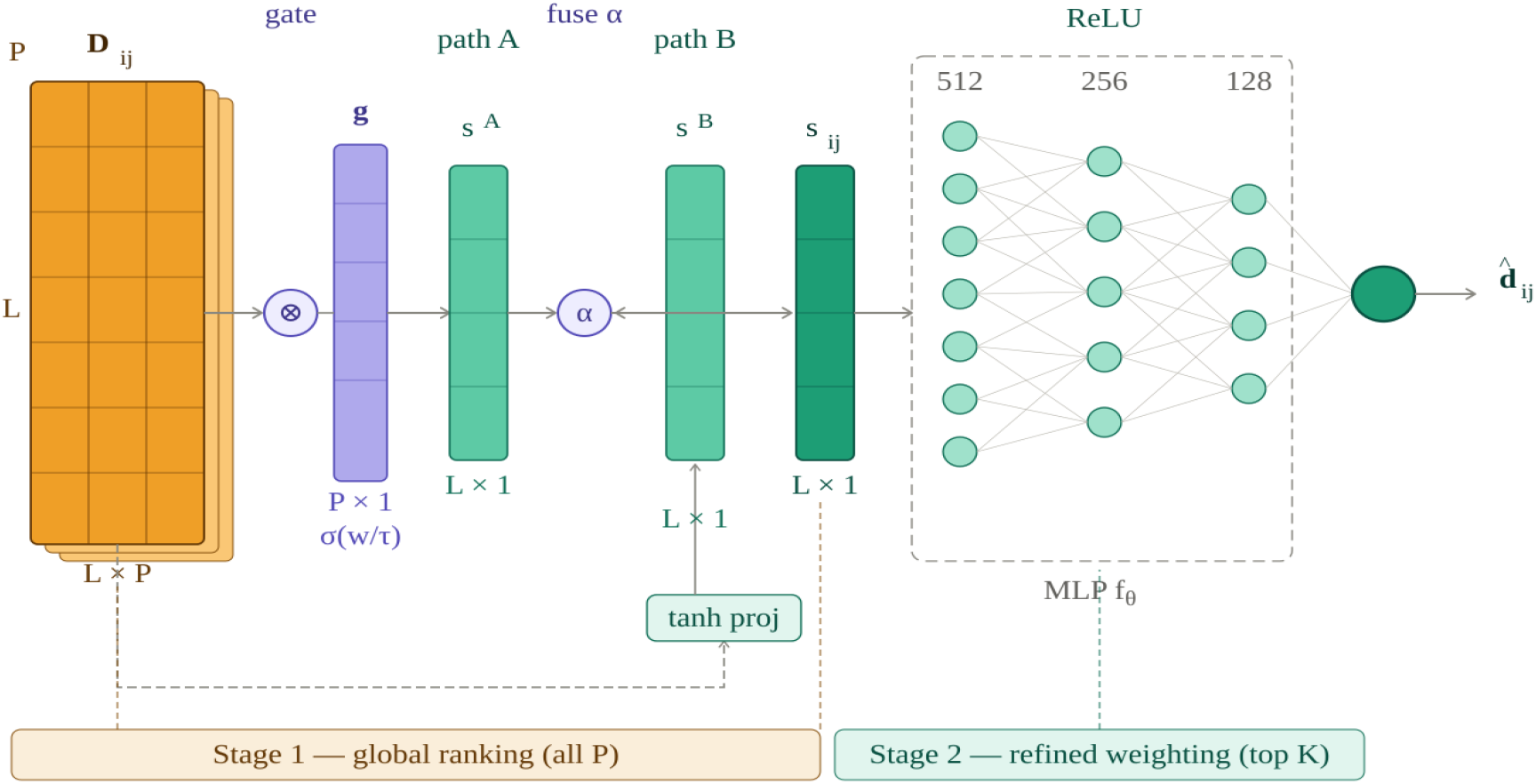
Architecture of the Differentiable Property Ranking Network (DPRN).

#### 4) Two-Path Sequence Compression

Given the descriptor matrix *D*_*ij*_ ∈ ℝ^*L×P*^ for a sequence pair (*s*_*i*_, *s*_*j*_), the DPRN compresses it into a position-level summary vector *S* _*ij*_ ∈ ℝ^*L*^ through two complementary paths that are fused via a learnable scalar *α* ∈ (0, 1):

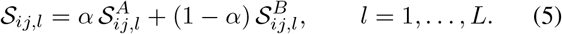

Here, Path A computes a gate-weighted linear sum of property differences at each position, providing an interpretable additive summary, while Path B applies a small position-shared nonlinear projection to capture non-additive interactions among properties. The full formulations of the gate function, the two paths, and the fusion mechanism are provided in Appendix C.

#### 5) Regression and Training Objective

The fused vector *S*_*ij*_ is then mapped to a predicted antigenic distance by a multilayer perceptron regressor *f*_*θ*_ : ℝ^*L*^ → ℝ, with 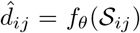 All learnable parameters — gates, temperature, projection weights, fusion scalar, and MLP weights — are trained jointly by minimising a mean squared error loss with an L1 penalty on the gates, which encourages uninformative properties to be suppressed. Full details of the loss function and optimisation are given in Appendix C.

#### 6) Two-Stage Property Selection

To improve stability, property selection is carried out in two stages. At Stage 1, the DPRN is trained on the full set *P* , and the *K* properties with the highest gate values form the reduced set *P*^*′*^. At Stage 2, a fresh DPRN is trained from scratch on *P*^*′*^ with a relaxed L1 coefficient, and the converged gate values are normalised to yield the final property importance weights { *w*^(*p*)^} _*p* ∈ *P*_*′* . The two-stage procedure is applied independently to scalar indices and substitution matrices, producing a separate top-*K* subset and weight vector for each category. The detailed two-stage equations are given in Appendix C.

### C. Input Feature Vector Construction

After the effective property subsets and their associated weights are identified through the property selection procedure, a unified input feature vector is constructed for each viral sequence pair (*s*_*i*_, *s*_*j*_). This vector jointly encodes both the positional sequence differences between the two strains and their physicochemical property representations, and serves as the direct input to the downstream prediction model. We follow the procedure described in [27] for constructing the input feature vector.

For a given sequence pair (*s*_*i*_, *s*_*j*_) of length *L*, the *binary difference vector* **b**_*ij*_ ∈ {0, 1}^*L*^ encodes position-wise sequence differences between the two strains. Each entry indicates whether the amino acids at position *l* differ between strains *s*_*i*_ and *s*_*j*_, as defined by the indicator function:

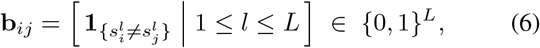

The binary difference vector **b**_*ij*_ thus provides a sparse, position-resolved summary of the mutational divergence between two Influenza A strains.

Let 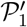 and 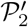 denote the two selected property subsets obtained from the property selection procedure, with associated optimized weight vectors 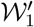 and 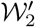 , respectively.

For each sequence pair (*s*_*i*_, *s*_*j*_), property descriptor matrices 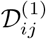 and 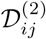 are generated from 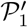 and 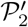 , encoding the physicochemical properties of the two strains at each sequence position. The combined representations 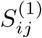 and 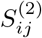 for the sequence pair (*s*_*i*_, *s*_*j*_) are then obtained by projecting each property descriptor matrix through its corresponding weight vector:

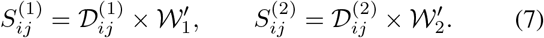

The resulting combined representations 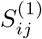and 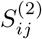 provide compact, property-weighted encodings of the two strains that capture both the identity and the weighted contribution of each selected property.

The final input feature vector *X*_*ij*_ for the sequence pair (*s*_*i*_, *s*_*j*_) is formed by concatenating the binary difference vector **b**_*ij*_ with the two property-weighted fused representations 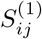 and 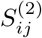:

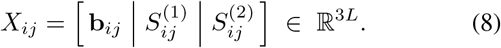

The dimensionality 3*L* arises from the concatenation of three *L*-dimensional vectors, where *L* is the length of the HA1 region of the viral protein sequence (approximately 330). Now, the final training set is defined as T_train_ = {(*X*_*ij*_, *d*_*ij*_) | (*s*_*i*_, *s*_*j*_) ∈ *S*_train_}.

### D. Dimensionality Reduction

The raw feature vectors representing each pair of Influenza A strains have dimension *M* = 3*L* ≈ 990. This dimensionality is modest by the standards of classical machine learning and would also be tractable for qubit-efficient quantum encodings such as amplitude encoding, which can, in principle, embed *M* real features into only ⌈log_2_ *M*⌉ qubits. In this work, however, we deliberately adopt an *angle-encoding* scheme, in which each classical value is loaded as a rotation angle of an *R*_*Z*_ or *R*_*X*_ gate. Because the number of qubits is strictly limited on both current quantum simulators and real quantum hardware, and because the circuit width is fixed in advance in our experiments, the encoding capacity of the circuit is far smaller than *M* . It is therefore the combination of the chosen angle-encoding scheme and the constrained qubit budget — rather than any intrinsic high-dimensionality of the data — that makes a feature vector of dimension *M* ≈ 990 incompatible with direct quantum encoding. A dimensionality reduction step is consequently essential: it compresses the original feature space of dimension *M* = 3*L* into a low-dimensional representation of dimension:

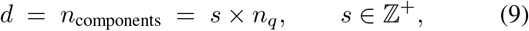

where *n*_*q*_ is the number of qubits and *s* is an integer scaling factor that sets how many input components are encoded per qubit. A larger *s* increases the information capacity of the reduced representation (and hence the number of rotation angles loaded into the circuit) at the cost of greater circuit depth; *s* is therefore treated as a hyperparameter that trades representational richness against circuit complexity.

A supervised Multi-Layer Perceptron (MLP) encoder was implemented as the reduction method, learning a nonlinear end-to-end mapping *f*_***θ***_ : ℝ^*M*^ →ℝ^*d*^ that directly minimises a combination of prediction error and, optionally, reconstruction error. A scaled hyperbolic-tangent transformation at the output bounds the reduced representation to [ − *π*, +*π*], ensuring valid rotation angles for the downstream quantum circuit.

#### Architecture

As illustrated in Fig. 3, the network comprises three components: an encoder body, a prediction head, and a decoder head used only during training. The encoder stacks residual blocks (pre-activation layer normalisation, linear transformation, ReLU activation, and dropout) followed by a linear bottleneck layer that produces the *d*-dimensional representation. The prediction head maps the bottleneck to the scalar antigenic distance estimate *ŷ*, while the optional decoder head mirrors the encoder in reverse to reconstruct the standardised input 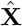 and is discarded at inference time.

**Fig. 3.**
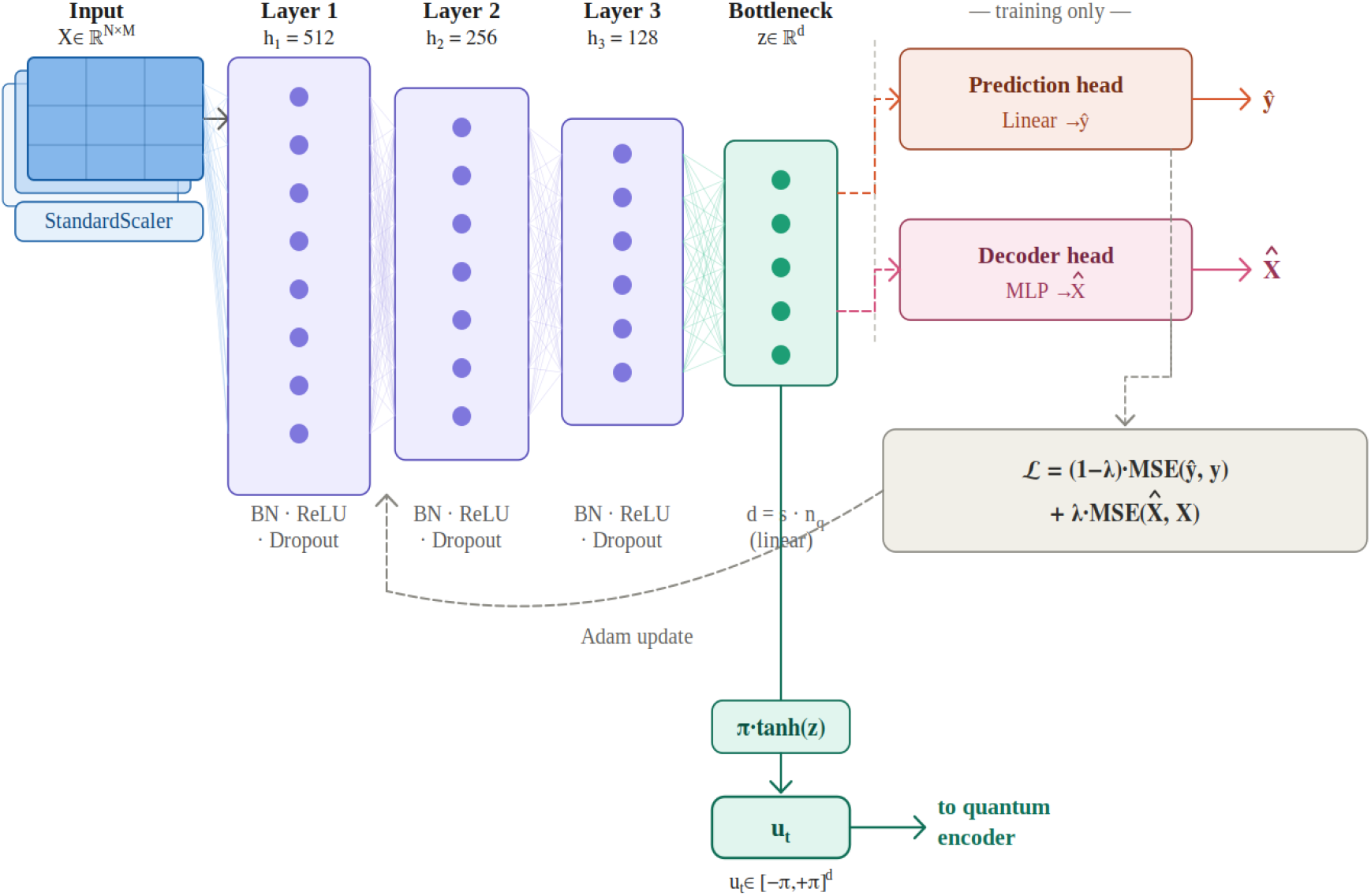
Supervised Neural Network Encoder (NNEncoder) Architecture for Dimensionality Reduction.

#### Training

Inputs and targets are standardised to zero mean and unit variance, with targets additionally clamped to suppress outlier pairs. The encoder is trained end-to-end by minimising the composite loss:

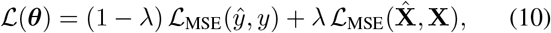

where *λ* ∈ [0, 1] controls the weight of the optional reconstruction term.

#### Encoder output

At inference time, only the encoder body and bottleneck layer are retained. The bottleneck output **z** is passed through a scaled hyperbolic tangent to produce valid quantum rotation angles:

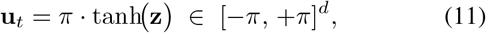

ensuring every component of **u**_*t*_ lies within the valid range of the *R*_*Z*_ and *R*_*X*_ rotation gates used in the downstream quantum encoding circuit.

### E. Hybrid Quantum-Classical Reservoir Computing

The next stage of the proposed framework implements a Hybrid Quantum-Classical Reservoir Computing (hQCRC) architecture, inspired by the design introduced in [37]. The pipeline consists of three sequential phases: (i) quantum data encoding, (ii) classical reservoir state update, and (iii) linear readout training via ridge regression. A schematic overview of the full hQCRC workflow is depicted in Fig. 4.

**Fig. 4.**
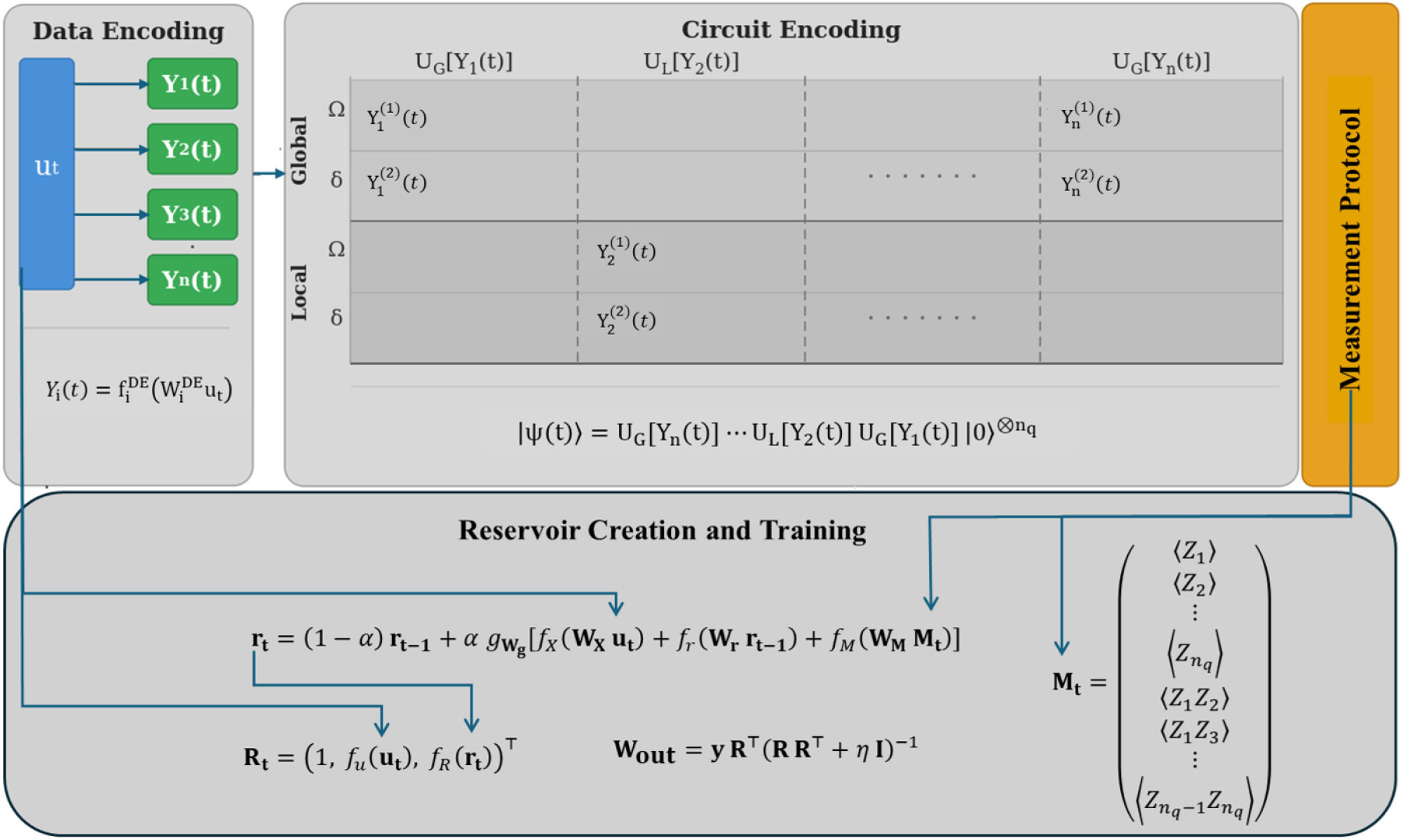
Architecture of hybrid Quantum-Classical Reservoir Computing (hQCRC).

#### 1) Phase 1: Quantum Data Encoding

The goal of the quantum encoding phase is to project the dimensionality-reduced input vector **u**_*t*_ ∈ [− *π*, +*π*]^*d*^, produced by the NNEncoder of Section 3.4, into a high-dimensional quantum Hilbert space *ℋ*, thereby generating a rich, non-linearly transformed measurement vector **M**_*t*_ for consumption by the classical reservoir. This phase is itself decomposed into two sub-steps: classical data encoding and parameterised quantum circuit encoding.

##### a) Step 1a-Classical Data Encoding

Before **u**_*t*_ is loaded into the quantum circuit, it is first projected into *n*_layers_ independent encoding vectors, one per circuit layer. For each layer *i* = 1, … , *n*_layers_, a fixed random projection matrix 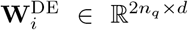 and a layer-specific nonlinear activation function 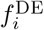 are applied to produce the encoded input for that layer:

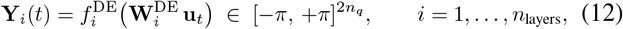

where the output dimension 2*n*_*q*_ supplies exactly two rotation angles per qubit per layer (one for *R*_*Z*_ and one for *R*_*X*_), and the output is clipped to [ -*π*, +*π*] to ensure validity as quantum rotation angles. The matrices 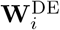 are drawn once from a uniform distribution at initialization and kept fixed throughout training, consistent with the reservoir computing principle of untrained random projections. The activation functions 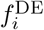 cycle through the set { tanh, sin, cos, id across layers,} introducing diverse nonlinear transformations of the input.

### b) Step 1b-Quantum Circuit Encoding

The encoded vectors 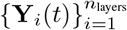are used as rotation angles in a parameterised quantum circuit inspired by the Transverse-Field Ising Model (TFIM) [38]. The circuit acts on *n*_*q*_ qubits, all initialised in the even superposition state | + *⟩* ^*⊗n*^*q* via Hadamard gates. A central design feature of the circuit is the distinction between two types of data-encoding layers that alternate throughout the circuit: *global* layers *U*_*G*_ and *local* layers *U*_*L*_.

- **Global layer** *U*_*G*_[**Y**_*i*_(*t*)] (odd-indexed layers *i* = 1, 3, 5, …): a single pair of angles (*δ*_*i*_, Ω_*i*_) is applied *identically to every qubit simultaneously*. In gate-model terms, this corresponds to the same *R*_*Z*_ and *R*_*X*_ rotation being broadcast across the full register:

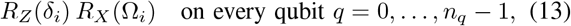

where 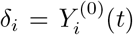 and 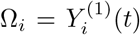are the first two components of **Y**_*i*_(*t*). Because all qubits receive the same angles, only 2 parameters are consumed from the encoded vector, regardless of the number of qubits.
- **Local layer** *U*_*L*_[**Y**_*i*_(*t*)] (even-indexed layers *i* = 2, 4, 6, …): each qubit *q* receives its own *independent* pair of angles (*δ*_*i,q*_, Ω_*i,q*_), enabling a richer, qubit-specific data encoding:

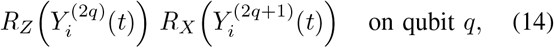

for *q* = 0, … , *n*_*q*_ −1. In this case 2*n*_*q*_ parameters are consumed from **Y**_*i*_(*t*), one pair per qubit. This provides significantly greater expressivity by allowing each qubit to be driven with a different amplitude and detuning control parameters.

Following the data-encoding sublayer (global or local), every layer *i* applies the same two fixed operations:

#### 1) Ising entangling sublayer

for each pair of qubits (*q*_*a*_, *q*_*b*_) with *q*_*a*_ < *q*_*b*_, apply an *R*_*ZZ*_ two-qubit gate with a fixed (untrained) coupling *J*_*q*_*a*_*q*_*b* drawn once at initialisation from U(−*κ*, +*κ*):

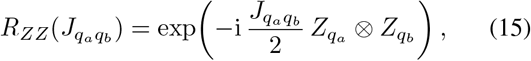

where *κ* is the coupling-strength hyperparameter. These fixed couplings generate entanglement and serve as the quantum reservoir weights—they are set at initialization and never updated during training.

#### 2) Global transverse-field pulse

a single *R*_*X*_ (Ω) rotation with a fixed global amplitude hyperparameter Ω is applied uniformly to all qubits: *R*_*X*_ (Ω)^*⊗n*^*q* . This pulse mixes qubit states after each entangling block and is always global, independently of whether the data-encoding sublayer of the same layer is global or local.

After all *n*_layers_ layers have been applied, the full quantum state takes the form

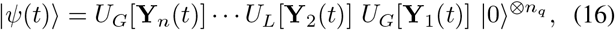

where the operators alternate between global (*U*_*G*_, odd layers) and local (*U*_*L*_, even layers) unitaries. The alternating global– local pattern maximizes the diversity of the encoded input across layers: global layers provide a computationally cheap, hardware-efficient broadcast encoding, while local layers introduce qubit-specific variation that increases the expressivity of the quantum feature map.

#### c) Step 1c: Measurement and Measurement Vector

Classical information is extracted from ⟨*ψ*(*t*)⟩ by computing the expectation values of a set of Pauli observables. Specifically, the measurement vector **M**_*t*_ is constructed from single-qubit *Z* expectation values and all pairwise *ZZ* two-qubit correlators:

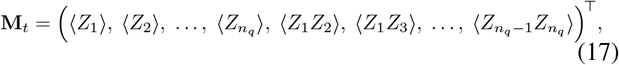

where each expectation value is evaluated as ⟨*O*⟩ = ⟨*ψ*(*t*)|*O*|*ψ*(*t*)⟩ ∈ [−1, 1].

#### 2) Phase 2: Classical Reservoir State Update

The measurement vector **M**_*t*_ is fed into a hybrid Quantum-Classical ESN reservoir [28, 39, 40], which is a specific kind of Recurrent Neural Network (RNN), that integrates quantum measurements with a recurrent classical state, providing the fading memory that pure quantum systems cannot sustain due to the destructive nature of projective measurements.

#### a )Reservoir state update

At each time step *t*, the classical reservoir state **r**_*t*_ ∈ ℝ^*N*^*r* is updated via the leaky integration rule:

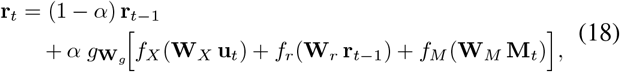

where *α* ∈ (0, 1] is the leak rate controlling the time-scale of the reservoir memory; **W**_*X*_ ∈ ℝ^*N*^*r* ^*×d*^, **W**_*r*_ ∈ ℝ^*N*^*r* ^*×N*^*r* , and **W**_*M*_ ∈ ℝ^*N*^*r* ^*×D*^ are fixed random weight matrices for the input, recurrent, and measurement contributions, respectively; *f*_*X*_ , *f*_*r*_, and *f*_*M*_ are element-wise nonlinear activation functions (implemented as tanh throughout); and *g*_**W**_*g* [·] ≡ tanh(**W**_*g*_ ·) is an additional nonlinear mixing layer with fixed random matrix **W**_*g*_ ∈ ℝ^*N*^*r* ^*×N*^*r* . Setting *f*_*M*_ ≡ **0** (i.e. **M**_*t*_ = **0**) recovers the classical-only ESN, enabling controlled ablation of the quantum contribution.

### b) Reservoir vector

After updating **r**_*t*_, the full reservoir vector **R**_*t*_ is assembled as:

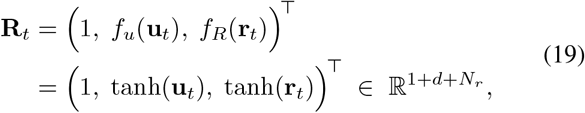

where the leading constant 1 serves as a bias term, *f*_*u*_ and *f*_*R*_ are element-wise tanh activations applied to the reduced input and reservoir state, respectively. The reservoir vectors for all training samples are collected into the matrix **R** = [**R**_1_, **R**_2_, … , **R**_*N*_]^*⊤*^ ∈ ℝ^*N ×*(1+*d*+*N*^*r*).

In this work, antigenic distance prediction treats each strain pair as an independent sample (i.e. there are no temporal dependencies across rows), so the reservoir state is reset to **r**_0_ = **0** before processing each sample. Under this condition the recurrent term *f*_*r*_(**W**_*r*_**r**_*t−*1_) vanishes, allowing the entire training batch to be processed with three parallelised matrix multiplications, which significantly reduces computation time for large datasets.

#### 3) Phase 3: Linear Readout via Ridge Regression

The final phase trains a linear output layer **W**_out_ that maps reservoir vectors **R**_*t*_ to antigenic distance predictions. The output weights are obtained in closed form via ridge regression:

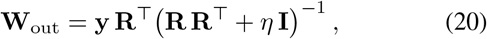

where **y** ∈ ℝ^*N*^ is the vector of training targets (antigenic distances), and *η >* 0 is the ridge regularisation coefficient that stabilises the matrix inversion and controls overfitting. Because the bias 1 is already embedded in **R**_*t*_, no intercept term is fitted separately. At inference time, the predicted antigenic distance for a new sample with reservoir vector **R**_*t*_ is: *ŷ*_*t*_ = **W**_out_ **R**_*t*_.

The key computational advantage of this formulation is that the only trainable component of the entire hQCRC pipeline is **W**_out_, which has a direct closed-form solution and requires no iterative gradient-based optimisation. All other components the data encoder projections 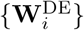 , the quantum circuit couplings { *J*_*ij}*_ , and the classical reservoir matrices **W**_*X*_ , **W**_*r*_, **W**_*M*_ , **W**_*g*_ — are fixed random matrices drawn once at initialisation and never updated.

## IV. Experiments

### A. Experimental Setup

Two datasets are used in the evaluations, covering the periods 1963–2003 and 2003–2025. For each dataset, samples are randomly partitioned into training (S_train_, 70%), validation (S_valid_, 10%), and test (S_test_, 20%) sets. Because the property selection procedure is supervised by the antigenic distances *d*_*ij*_, applying it to the full dataset would risk information leakage and inflated test performance. To prevent this, the test pairs were held out before Stage 1, and all supervised components — including property weighting, top-ranked property selection, weight refinement, model selection, and final regressor training — were carried out only on the training/validation data. The test pairs and their antigenic distances were untouched until the final evaluation, so the reported test *R*^2^, MAE, and RMSE genuinely measure generalisation to sequence pairs unseen by both the feature-selection procedure and the prediction model.

At the Property Selection stage, the top *K* = 5 subsets and their associated weight vectors are retained for each property category. This value was chosen to balance expressiveness— preserving sufficient biochemical diversity across the selected properties—against the risk of overfitting introduced by weakly relevant or redundant features. The DPRN model is trained using the AdamW optimiser [41] with an initial learning rate of 10^*−*3^ and a weight decay of 10^*−*4^.

State-vector simulation scales exponentially in the number of qubits *n*_*q*_, which makes the circuit width the primary computational bottleneck of our pipeline. While simulators can in principle support up to approximately 30 qubits and current neutral-atom hardware up to approximately 156 qubits, circuits with more than *n*_*q*_ = 14 qubits become computationally prohibitive in practice, with full training runs exceeding a day. We therefore fix *n*_*q*_ = 14 throughout all experiments, and the remaining encoding hyperparameters are configured relative to this fixed qubit budget. Within this constraint, the scaling factor *s* at the dimensionality reduction stage controls the information capacity of the bottleneck representation relative to the qubit count. We select *s* by cross-validation over the candidate set *s* ∈ { 8, … , 15}, treating it as a hyperparameter jointly with *n*_qubits_ and *n*_layers_. For the small dataset (1963– 2002), *s* = 8 yielded the best performance and was adopted as the primary configuration, whereas for the large dataset (2003– 2025), *s* = 15 was used instead. The neural network encoder at this stage is a four-layer MLP with hidden dimensions [1024, 512, 256, 128], trained with a learning rate of 5 × 10^*−*4^ and optimised using the Adam algorithm [42].

The hQCRC framework involves hyperparameters belonging to three distinct phases: Quantum Data Encoding, Classical Reservoir, and Linear Readout. Table I summarises the key hyperparameters and their default values used in this study. The quantum encoding circuit instantiated on *n*_*q*_ = 14 qubits with *n*_layers_ = 3 encoding layers. Because the same fixed circuit is evaluated independently for every sample, the encoding of the full dataset is embarrassingly parallel; The resulting measurement vectors are then concatenated. Classical information is extracted from the evolved quantum state by estimating the expectation values of a fixed set of Pauli-*Z* observables. The measurement vector **M**_*t*_ comprises all singlequbit expectation values *⟨Z*_*i*_⟩ and all two-qubit correlators ⟨*Z*_*i*_*Z*_*j*_⟩ over every distinct qubit pair. Three-body ⟨*Z*_*i*_*Z*_*j*_*Z*_*k*_⟩ correlators were not included. All expectation values are computed using the Qiskit `AerEstimatorV2`, which provides numerically exact expectation values and avoids the statistical noise that finite-shot sampling would introduce, ensuring that the reported results isolate the effect of the encoding rather than measurement noise.

**TABLE I.** Key hyperparameters of hQCRC pipeline and default values.

| Phase | Hyperparameter | Symbol | Default |
| --- | --- | --- | --- |
| Quantum data encoding | Number of qubits | $n_q$ | 14 |
| | Number of circuit layers | $n_{\text{layers}}$ | 3 |
| | Coupling strength | $\kappa$ | 3.0 |
| | Global transverse field | $\Omega$ | 0.3 |
|  | Include ZZZ correlators | — | False |
| Classical reservoir | Reservoir size | $N_r$ | 2048 |
| | Leak rate | $\alpha$ | 0.3 |
| | Spectral radius | $\rho$ | 0.9 |
| Linear readout | Ridge regularisation | $\eta$ | $10^{-4}$ |

### B. Evaluation Metrics

Model performance is assessed using three complementary regression metrics evaluated on the held-out test set S_test_. Throughout, *d*_*ij*_ and 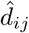 denote the true and predicted antigenic distances for the strain pair (*s*_*i*_, *s*_*j*_), respectively, and *N*_*Test*_ = |S_test_| is the number of test samples.

#### 1) Mean Absolute Error (MAE)

MAE quantifies the average magnitude of prediction errors:

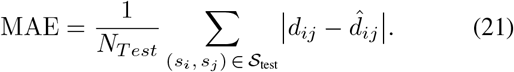

In the context of antigenic distance prediction, MAE directly quantifies how far, on average, a predicted antigenic distance deviates from the experimentally derived value. Because each residual is weighted equally, MAE provides a straightforward and interpretable measure of the average deviation between predictions and true values.

#### 2) Root Mean Squared Error (RMSE)

RMSE penalises large errors more heavily than MAE owing to the squared residuals:

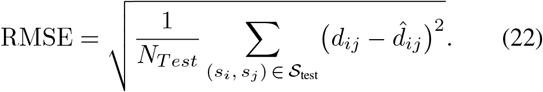

RMSE measures the typical size of the prediction error while giving greater influence to strain pairs for which the predicted antigenic distance is substantially different from the true distance. This metric is therefore more sensitive to outliers and systematic deviations, making it a useful complement to MAE.

#### 3) Coefficient of Determination (R^2^)

*R*^2^ measures the proportion of variance in the true distances that is explained by the model:

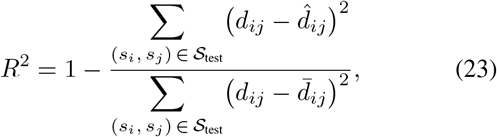

where 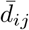 denotes the mean of the true antigenic distances over the test set. *R*^2^ thus measures how well the model captures the overall variation in antigenic distances across strain pairs, rather than only the average size of individual errors.

### C. Baselines

Our comparative analysis is conducted to demonstrate the effectiveness of the proposed model for antigenic distance prediction against several representative baseline methods. Specifically, we compare with four established A/H3N2 prediction models: (1) Durazzi et al. [22] adopted a sequence-based approach using BiLSTM embeddings combined with ridge regression, without explicitly using amino acid property information. (2) Yao et al. [18] employed a random forest regression model to select informative substitution matrices from AAindex2, capturing antigenicity-related patterns. (3) Li et al. [25] used manually engineered antigenic features together with XGBoost regression to achieve strong predictive performance.

(4) Finally, Geng et al. [27] introduced an attention-based framework that extracts and weights antigenicity-related features from AAindex1 and AAindex2, followed by a multilayer perceptron for prediction. The latter three methods explicitly incorporated amino acid properties into the model inputs.

## V. Results and Discussion

### A. Property Selection

To effectively select informative features, we apply a two-stage property selection procedure separately to scalar physicochemical indices and substitution matrices for both small (1963-2002) and large (2003–2025) datasets. The selected properties and their normalized contributions illustrate the most informative descriptors for antigenicity prediction. Fig. 5 shows the top 5 selected Amino Acid properties and their normalized weights that belong to the large dataset.

**Fig. 5.**
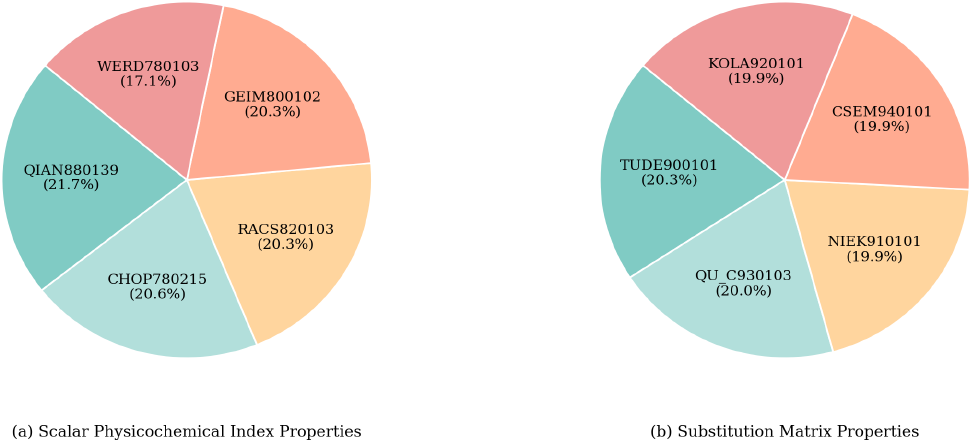
Selected Amino Acid Properties and Normalized Weights from the 2003–2025 Dataset.

### B. Performance Comparison

In this section, FluQRC is compared with four baseline methods on both datasets, and the results are presented in Table II. All baselines were re-evaluated under the same unified experimental setting used for FluQRC, so that the metrics are directly comparable across methods.

**TABLE II.** Model Performance Comparison on the 1963–2002 and 2003–2025 datasets.

| Study | Year | MAE | | RMSE | | $R^2$ | |
| --- | --- | --- | --- | --- | --- | --- | --- |
|  |  | 1963–2002 | 2003–2025 | 1963–2002 | 2003–2025 | 1963–2002 | 2003–2025 |
| Durazzi et al. [22] | 2025 | 1.114 | 1.291 | 1.604 | 1.723 | 0.886 | 0.211 |
| Yao et al. [18] | 2017 | 0.902 | 1.292 | 1.153 | 1.661 | 0.961 | 0.554 |
| Li et al. [25] | 2024 | 0.581 | 0.798 | 0.783 | 1.063 | 0.981 | 0.722 |
| Geng et al. [27] | 2025 | 0.544 | 0.463 | 0.752 | 0.741 | 0.980 | 0.862 |
| **FluQRC (Ours)** | **2026** | **0.545** | **0.369** | **0.734** | **0.635** | **0.982** | **0.900** |

#### Effect of amino acid property integration

In both datasets, the methods that explicitly incorporate amino acid properties into the model inputs—FluQRC, Geng et al., Li et al., and Yao et al.— achieve lower MAE and RMSE as well as higher *R*^2^ than Durazzi et al., which relies solely on protein language-model embeddings without incorporating amino acid property information. This indicates that although language-model embeddings are effective for broad antigenic cluster prediction, they may overlook important site-specific physicochemical details.

#### Comparison among property-aware methods

Among the methods that explicitly incorporate amino acid properties, Li et al. [25] proposed a comprehensive set of twelve handcrafted features derived from the HA sequence fed into an XGBoost regression model. This richer feature representation is the main reason Li et al. outperforms Yao et al. [18], whose Joint Random Forest Regression (JRFR) method restricts its search space to substitution matrices sourced exclusively from the AAIndex2 database, an inherently narrower information source that limits the diversity of learnable features. Geng et al. [27] (FluAttn) extend this idea further by leveraging both AAIndex1 and AAIndex2 through an attention-based feature mining framework that automatically selects and weights the most antigenicity-relevant properties, rather than relying on handcrafted feature engineering. The resulting diversity and breadth of the feature space—combined with a multi-head static attention mechanism that quantifies the differential contribution of each property—gives FluAttn a clear advantage over both Li et al. and Yao et al.

Our Framework (FluQRC) benefits from a pipeline that integrates three novel components: (i) a Differentiable Gated Property Ranking Network for data-driven property selection, a Dimensionality Reduction network that compresses the feature representation into a form suitable for quantum processing, and (iii) a hybrid Quantum-Classical Reservoir Computing architecture for antigenic distance regression. Together, these components allow FluQRC to mine antigenicity-relevant features from a rich and diverse set of information sources more efficiently than any of the baselines. As a result, on the small 1963–2002 dataset FluQRC achieves performance comparable to Li et al. and Geng et al., while on the larger and more challenging 2003–2025 dataset FluQRC substantially outperforms all baselines across all three metrics, demonstrating the scalability and effectiveness of the proposed architecture in large-scale and complex settings.

### C. Ablation Experiments: Classical Reservoir vs Hybrid Reservoir

To isolate the contribution of the quantum reservoir, we replaced it with a pure classical reservoir of equivalent size while keeping all other pipeline components identical. The results are reported in Table III.

**TABLE III.** Hybrid Quantum-Classical Reservoir vs Pure Classical Reservoir of equivalent size.

| Metric | Classical |  | Hybrid |  |
| --- | --- | --- | --- | --- |
|  | 1963–2002 | 2003–2025 | 1963–2002 | 2003–2025 |
| MAE | 0.562 | 0.381 | 0.545 | 0.369 |
| RMSE | 0.746 | 0.642 | 0.734 | 0.635 |
| $R^2$ | 0.982 | 0.897 | 0.982 | 0.900 |
| Exec. Time (min) | 2.3 | 16.4 | 5.2 | 39.1 |

The quantum reservoir yields modest but consistent improvements across all metrics and both datasets. On the 1963– 2002 dataset, MAE decreases by 3.02%, RMSE by 1.61%, and *R*^2^ remains unchanged. On the larger 2003–2025 dataset, MAE improves by 3.15%, RMSE by 1.09%, and *R*^2^ increases by 0.33%. Although the improvements are modest rather than dramatic; these results confirm that the hybrid quantum-classical reservoir provides a systematic predictive advantage over a classical reservoir of equivalent size, likely due to its ability to generate an entanglement-rich feature space that is not efficiently reproducible classically at the same scale.

This advantage comes with an additional computational cost. The quantum reservoir increases execution time by approximately 2.3 × on the small dataset and 2.4 × on the large dataset. Whether this trade-off is acceptable in practice depends on the deployment context: for offline batch screening of candidate vaccine strains, the additional time is unlikely to be a bottleneck, whereas for real-time surveillance applications it may warrant further optimisation of the encoding circuit. Taken together, this ablation confirms that the quantum reservoir is a beneficial component of FluQRC and is responsible for a consistent, if incremental, portion of the performance gains over the classical baseline.

## VI. Conclusion

We introduced FluQRC, a hybrid quantum-classical reservoir computing framework for predicting antigenic distance between influenza A/H3N2 strains from hemagglutinin sequence data. The framework combines three synergistic components. First, the Differentiable Gated Property Ranking Network (DPRN) automatically identifies and weights the most informative physicochemical and substitution-matrix descriptors in a fully data-driven manner, eliminating the need for manual feature engineering and substantially broadening the diversity of the biochemical information encoded into the model. Second, the supervised NNEncoder compresses the resulting high-dimensional pairwise feature vectors into compact, bounded representations that are directly compatible with quantum rotation gates. Third, the hQCRC architecture exploits the entanglement-rich feature space of a Transverse-Field Ising Model-inspired quantum circuit, augmented by a classical ESN, to generate expressive reservoir states from which antigenic distances are predicted by a computationally efficient closed-form linear readout.

Ablation experiments confirm that the quantum reservoir provides a consistent predictive advantage over a classical reservoir of equivalent size across both datasets, at the cost of a roughly 2.3–2.4 × increase in execution time. Comparison against four established baselines demonstrates that FluQRC achieves competitive performance on the small historical dataset (1963–2002) and substantially outperforms all baselines on the larger and more challenging 2003–2025 dataset, where it attains MAE = 0.369, RMSE = 0.635, and *R*^2^ = 0.900. These results confirm that the combination of automated property selection, neural dimensionality reduction, and hybrid quantum-classical reservoir computing scales effectively to large serological datasets.

Several directions remain open for future work. The quantum encoding circuit was simulated using exact statevector methods; deploying FluQRC on near-term quantum hardware will require investigating the impact of shot noise and device-specific noise models on prediction accuracy. Furthermore, extending the framework to additional influenza subtypes, such as H1N1 or avian strains, and evaluating cross-subtype generalisation via transfer or meta-learning strategies represents another natural avenue.

## Appendix

### A. Reservoir Computing

Reservoir Computing (RC) [28] is a supervised learning framework designed to efficiently process temporal or sequential data. The architecture typically consists of three main components: an input layer, a reservoir, and a readout layer, as depicted in Fig. 1. The central idea of RC is to project the input signal into a high-dimensional dynamical space through a fixed nonlinear system known as the reservoir. This transformation produces rich internal representations of the input sequence, enabling the readout layer to learn the desired mapping using relatively simple training procedures. RC systems can be implemented using either computational models such as recurrent neural networks or physical dynamical systems, including mechanical oscillators, optical systems, fluid dynamics, and quantum devices [43, 44].

A key requirement for a functional reservoir is the Echo State Property (ESP). This property ensures that the reservoir state gradually becomes independent of its initial conditions and depends primarily on recent inputs. In practical terms, ESP guarantees that the system possesses fading memory, meaning that the influence of earlier inputs diminishes over time while more recent inputs dominate the system dynamics [45]. The duration required for the reservoir to forget its initial state is commonly referred to as the washout period. One of the most well-known implementations of RC is the Echo State Network (ESN)^3^ [28, 39, 40], where the reservoir is realized as a large recurrent neural network (RNN) with randomly generated and fixed internal connections. Unlike traditional recurrent neural networks, the weights within the reservoir are not optimized during training. Instead, only the output layer parameters are learned. This design significantly reduces the computational cost associated with training, as expensive gradient-based optimization of the recurrent structure is avoided.

Let the input time series be denoted **u**_1_, … , **u**_*T*_ , where each **u**_*t*_ ∈ ℝ^*n*^. The reservoir evolution is governed by a deterministic map *R*, which in standard digital implementations is parameterised by a set of fixed random matrices denoted collectively by ***W*** . At each discrete time step *t*, the reservoir state **r**_*t*_ ∈ ℝ^*N*^ is updated according to the recursive relation:

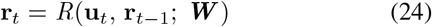

In standard digital implementations, *R* is realised as a leaky integrator update, which introduces a smoothing coefficient *α* ∈ [0, 1] to regulate the trade-off between the current input and the preceding reservoir state:

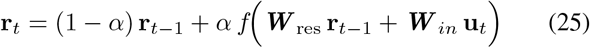

where ***W*** _in_ ∈ ℝ^*N ×n*^ and ***W*** _res_ ∈ ℝ^*N ×N*^ are the fixed random input and reservoir weight matrices, respectively; *f* is a pointwise nonlinear activation function; and *α* ∈ [0, 1] is the leaking rate, which regulates how strongly previous reservoir states influence the current state. For a suitable choice of *R*, the state **r**_*t*_ retains information about past inputs only up to a characteristic decay timescale known as the fading memory horizon.

The final component of the RC architecture is a linear readout layer parametrised by ***W*** _out_ ∈ ℝ^*m×N*^ . This layer is the sole trainable component of the system and maps the reservoir state to the predicted output **ŷ**_*t*_:

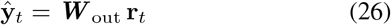

where **ŷ**_*t*_ ∈ ℝ^*m*^ is the network’s prediction at time step *t*. The readout matrix ***W*** _out_ is obtained as the closed-form solution to a Tikhonov-regularised (ridge regression) least-squares optimisation problem:

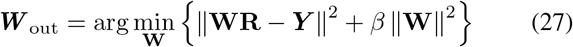

where **R** ∈ ℝ^*N ×T*^ collects the reservoir state vectors over the entire training sequence U; ***Y*** denotes the matrix of target output vectors; and *β* ∈ [0, 1] is the Tikhonov regularisation parameter that penalises large weight norms and prevents overfitting. The unique closed-form solution is given by:

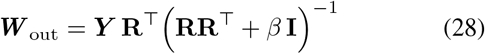

This closed-form solution requires only a single matrix inversion of order *N* × *N* and can be computed analytically without iterative gradient-based optimisation. The overall simplicity of the training procedure—updating solely ***W*** _out_ while keeping ***W*** _res_ and ***W*** _in_ frozen—represents one of the principal advantages of the RC paradigm over conventional RNNs, making it particularly attractive for resource-constrained and real-time learning applications.

#### B. Quantum Reservoir Computing

The estimation of expectation values in variational quantum algorithms, such as quantum neural networks, often suffers from the vanishing gradient problem, leading to nearly flat optimization landscapes known as barren plateaus barren plateaus [31]. This phenomenon significantly hinders the trainability of parameterized quantum circuits. As a promising alternative, quantum reservoir computing (QRC) [29, 30, 46] avoids gradient-based optimization altogether, thereby mitigating the challenges associated with barren plateaus.

QRC is a hybrid quantum–classical framework that builds upon the principles of classical reservoir computing, where a fixed high-dimensional dynamical system transforms input data into a rich feature space, and only a simple readout layer is trained. QRC adopts the same philosophy but replaces the classical reservoir with a quantum system, thereby leveraging the unique properties of quantum mechanics such as superposition, entanglement, and exponentially large Hilbert spaces.

The central idea of QRC is to use the Hilbert space of a quantum system as an enhanced feature space for input data. Because the dimension of the Hilbert space grows exponentially with the number of qubits, even small quantum systems can generate highly expressive representations. In this framework, input data are encoded into quantum states, and the intrinsic dynamics of the quantum system transform these inputs into complex, high-dimensional states. These states contain rich temporal and nonlinear correlations, making them well-suited for learning complex input–output relationships using simple classical models.

Formally, given a time-series input sequence *U* = {*u*_0_, *u*_1_, … , *u*_*t*_, … ,} the QRC process proceeds. At each time step *t* , the input *u*_t_ is first encoded into the quantum system via an encoding unitary operator *U*_*enc*_(*u*_*t*_), which maps classical data into the quantum state space. The system then evolves under a unitary transformation that governs the reservoir dynamics. In continuous-time formulations, this evolution is described by

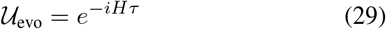

where ***H*** is the Hamiltonian of the quantum system and ***τ*** is the evolution time. This evolution spreads the encoded information across the system through quantum correlations and entanglement. In gate-based implementations, this evolution is typically approximated by a *θ*-parameterized quantum circuit, such that:

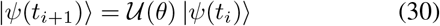

After evolution, the quantum state is measured to extract classical information. Specifically, expectation values of selected observables ***O*** (e.g., Pauli operators) are computed:

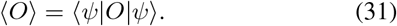

These measured values serve as features representing the transformed input data. The collection of these features across time constitutes the reservoir state, which is then fed into a classical machine learning model—typically a linear regressor or classifier—for training and prediction.

Overall, QRC forms a hybrid pipeline consisting of four main stages: (i) classical input encoding, (ii) quantum reservoir evolution, (iii) measurement-based feature extraction, and (iv) classical post-processing. A key advantage of this framework is that only the output layer requires training, while the quantum reservoir remains fixed, significantly reducing computational complexity. However, the effectiveness of QRC depends critically on the design of the quantum reservoir, as the circuit must be sufficiently complex to generate informative and diverse features that capture the underlying structure of the data.

Such architectures have demonstrated strong performance in learning complex temporal dependencies, particularly in time-series forecasting tasks [47, 48]. More generally, reservoir computing exploits the intrinsic dynamics of complex systems to project inputs into expressive representations, enabling efficient solutions for tasks such as prediction and classification [44, 49]. Although QRC was initially developed and widely applied for time-series analysis, its applicability is not limited to temporal data. Recent studies have shown that QRC can be effectively extended to a broader range of problems, including applications in quantum chemistry simulations and hybrid quantum–classical neural network design [32, 33]. These developments highlight that QRC serves as a versatile computational paradigm capable of addressing diverse machine learning and scientific computing tasks beyond its original scope.

#### C. Detailed Formulation of the Differentiable Gated Property Ranking Network

This appendix provides the full mathematical formulation of the Differentiable Property Ranking Network (DPRN) summarised in the main text. We describe (i) the learnable gate parameterisation, (ii) the two compression paths and their fusion, (iii) the regression and training objective, and (iv) the two-stage property selection procedure.

##### 1) Differentiable Gated Property Ranking Network

The central goal of property selection is to identify, from the full set P of descriptors, a compact subset P’^*′*^ of size *K* =|P’ | whose members are most informative for predicting antigenic distance. To achieve this, we propose a novel *Differentiable Property Ranking Network* (DPRN) — a neural architecture in which property importance is encoded directly as a set of learnable gate parameters that are optimized end-to-end alongside the prediction task. The design draws on the principle of differentiable feature selection through continuous relaxation of discrete gates [36], in which hard binary feature masks are replaced by smooth, differentiable surrogates that can be jointly trained with the downstream objective.

For each property *p* ∈ P , a raw logit *w*_*p*_ ∈ ℝ is introduced as a learnable scalar. The corresponding gate value is obtained by passing *w*_*p*_ through a sigmoid function controlled by a learnable temperature parameter *τ >* 0 [50]:

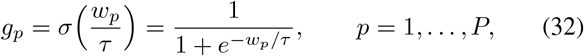

where *τ* = exp(log *τ*) is constrained to the interval (0.1, 10] via clamping, and log *τ* is jointly optimised during training. Crucially, each gate *g*_*p*_ is independent — unlike softmax normalisation, which forces competition among properties — allowing multiple properties to simultaneously receive high importance scores.

##### 2) Two-Path Sequence Compression

Given the descriptor matrix D_*ij*_ ∈ ℝ^*L×P*^ for a sequence pair (*s*_*i*_, *s*_*j*_), the DPRN compresses it into a position-level summary vector S_*ij*_ ∈ ℝ^*L*^ that encodes the gate-weighted property difference at each sequence position. Two complementary compression paths are computed and fused.

##### Path A – Gate-weighted sum

Each row of D_*ij*_ (corresponding to one sequence position) is contracted with the gate vector **g** = (*g*_1_, … , *g*_*P*_)^*⊤*^:

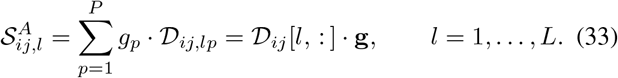

This path produces a linear summary of how much each position differs between the two sequences, weighting each property by its learned importance.

##### Path B – Shared nonlinear projection

A small two-layer network with weights shared across all sequence positions is applied independently at every position to produce a nonlinear summary:

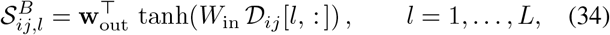

where *W*_in_ ∈ ℝ^*H*^*p*^*×P*^ and **w**_out_ ∈ ℝ^*H*^*p* are learnable parameters, and *H*_*p*_ = 32 in all experiments. Unlike Path A, Path B does not use the gate vector and is designed to capture non-additive interactions among properties at each site.

##### Fusion

The outputs of the two paths are combined via a learnable scalar *α* ∈ (0, 1):

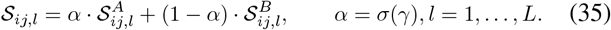

where *γ* ∈ ℝ is a learnable scalar and *σ*(·) denotes the sigmoid function. This yields the fused representation vector S_*ij*_ ∈ ℝ^*L*^, where each element S_*ij,l*_ reflects the net difference at position *l* under all properties, blending the interpretable linear weighting of Path A with the nonlinear correction of Path B.

##### 3) Regression and Training Objective

Given antigenic distances *d*_*ij*_, the training set is defined as T_train_ = {(*S*_*ij*_, *d*_*ij*_) | (*s*_*i*_, *s*_*j*_) ∈ S_train_}. A multilayer perceptron (MLP) regressor *f*_*θ*_ : ℝ^*L*^ → ℝ is trained on T_train_ to produce the predicted antigenic distance 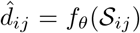 . The complete model ^*p*=1^gates 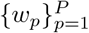 , temperature log *τ* , projection parameters {*W*_in_, **w**_out_}, fusion scalar *γ*, and MLP weights *θ* — is trained jointly by minimising a regularised mean squared error loss over the training set T_train_.

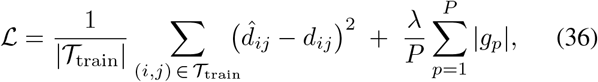

where *d*_*ij*_ is the observed antigenic distance and *λ* is the L1 sparsity coefficient, which encourages uninformative properties to have gates close to zero.

##### 4) Two-Stage Property Selection

A two-stage selection strategy is employed to separate the coarse identification of relevant properties from the fine-grained estimation of their relative weights. Directly learning weights over the full descriptor set *P* is susceptible to instability, as redundant or noisy properties can decrease the contributions of informative ones. The two stages address this by first narrowing the candidate set and then refining the weight distribution within it.

At stage 1, the DPRN is trained on the full property set P. Upon convergence, the gate values 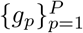 reflect the relative importance of each property as learned from the data. The *K* properties with the highest gate values are selected to form the reduced set P^*′*^:

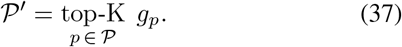

At Stage 2, a new DPRN instance is trained from scratch on the reduced property set P^*′*^ only, with the L1 regularisation coefficient reduced to *λ/*10 to allow finer gradient-driven differentiation among the retained properties. The gate values obtained at convergence in Stage 2 define the final normalised property importance weights:

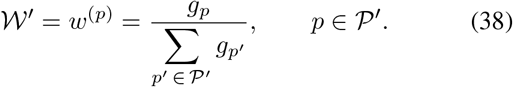

These weights {*w*^(*p*)^} _*p* ∈ *P*_*′* are used in the downstream antigenic distance predictor to combine the selected property descriptors into the input feature vector. The two-stage procedure is applied independently to each category of descriptors (scalar indices and substitution matrices), yielding separate top-*K* subsets and weight vectors for each category.

## Footnotes

1 https://www.gisaid.org

2 https://www.ncbi.nlm.nih.gov/genomes/FLU/

3 A specific kind of RNN designed to efficiently handle sequential data.

